# Relative performance of Oxford Nanopore MinION vs. Pacific Biosciences Sequel third-generation sequencing platforms in identification of agricultural and forest pathogens

**DOI:** 10.1101/592972

**Authors:** Kaire Loit, Kalev Adamson, Mohammad Bahram, Rasmus Puusepp, Sten Anslan, Riinu Kiiker, Rein Drenkhan, Leho Tedersoo

## Abstract

Culture-based molecular characterization methods have revolutionized detection of pathogens, yet these methods are either slow or imprecise. The second-generation sequencing tools have much improved precision and sensitivity of detection, but the analysis processes are costly and take several days. Of third-generation techniques, the portable Oxford Nanopore MinION device has received much attention because of its small size and possibility of rapid analysis at reasonable cost. Here, we compare the relative performance of two third-generation sequencing instruments, MinION and Pacific Biosciences Sequel in identification and diagnostics of pathogens from conifer needles and potato leaves and tubers. We demonstrate that Sequel is efficient in metabarcoding of complex samples, whereas MinION is not suited for this purpose due to the high error rate and multiple biases. However, we find that MinION can be utilized for rapid and accurate identification of dominant pathogenic organisms from plant tissues following both amplicon-based and metagenomics-based approaches. Using the PCR-free approach with shortened extraction and incubation times, we performed the entire MinION workflow from sample preparation through DNA extraction, sequencing, bioinformatics and interpretation in two and half hours. We advocate the use of MinION for rapid diagnostics of pathogens, but care needs to be taken to control or account for all potential technical biases.

**IMPORTANCE:** We develop new and rapid protocols for MinION-based third-generation diagnostics of plant pathogens that greatly improves the speed and precision of diagnostics. Due to high error rate and technical biases in MinION, PacBio Sequel platform is more useful for amplicon-based metabarcoding from complex biological samples.

## INTRODUCTION

Fungal pathogens and pests cause enormous losses in agriculture and forestry. Rapid and precise identification of pathogens enables efficient countermeasures and reduces the costs of biocides and losses to disease (Comtet et al., 2015). Direct morphology-based and culture-based diagnoses are often too imprecise or slow. Molecular methods such as specific primers, probes and sequence analysis are more accurate and can be rapidly applied to infected tissues (Kashyap et al., 2017). Although specific probes or primers combined with PCR/qPCR can be rapidly applied to tissue samples and environmental material, these methods lack the capacity to detect species or strains other than those intended, or worse, yield false positive signals (Grosdidier et al., 2017). Although high-quality sequencing reads are highly precise, Sanger sequencing of PCR products takes 1-3 days, depending on access to sequencing laboratory, and it may fail when DNA of several species or polymorphic alleles are amplified (Hyde et al., 2013).

Second- and third-generation high-throughput sequencing (HTS) platforms read hundreds of thousands to billions of DNA molecules, recovering the targeted taxa when present at very low proportions (Bik et al., 2012; Nilsson et al., 2019). However, library preparation and running of HTS instruments typically takes several days and there are queues of weeks to months in commercial service providers. Furthermore, a single run costs >500 EUR, which renders it unfeasible for rapid identification of pathogens (Tedersoo et al., 2019). In spite of millions of output reads, the second-generation SOLiD, Roche 454, Illumina and Ion Torrent platforms suffer from short sequence length that is suboptimal for accurate identification of microorganisms because of low taxonomic resolution of short marker gene fragments (100-500 bp; Mosher et al., 2014; Schloss et al., 2016). Third-generation sequencing platforms Pacific Biosciences (PacBio; RSII and Sequel instruments; www.pacb.com) and Oxford Nanopore (MinION, GridION and PromethION instruments; https://nanoporetech.com/) enable average sequence length of >20,000 bases, but this comes at 5%-20% error rate (Weirather et al., 2017; Jain et al., 2018; Tedersoo et al., 2018, 2019). In PacBio instruments, the built-in circular consensus sequencing generates multiple copies of the same fragment with highly accurate consensus (Eid et al., 2009; Rhoads and Au, 2015). Therefore, long consensus molecules have been readily used in de novo assembly of complex genomes (Giordano et al., 2017) and DNA barcoding (Hebert et al., 2018). PacBio-based metabarcoding analyses provide greater resolution compared with second-generation HTS tools in bacteria (Singer et al., 2016; Wagner et al., 2016; Schloss et al., 2016) and fungi (Tedersoo et al., 2018), including plant pathogens (Walder et al., 2017).

Compared with other HTS platforms represented by large and quite expensive machines, the Oxford Nanopore MinION device is of the size of a cell phone that costs ca. 900 EUR, making it affordable to governmental institutions, research laboratories and small companies (Mikheyev and Tin, 2014; Lu et al., 2016). Its small size and low power consumption enable carrying the device, basic analysis toolkit, batteries and computer virtually anywhere, as demonstrated by in situ runs in a tropical rain forest (Quick et al., 2016), Antarctic desert (Johnson et al., 2017) and space station (Castro-Wallace et al., 2017). MinION has a capacity to produce >1 million sequences per day, with maximum read length approaching 1,000,000 bases (Jain et al., 2018). Because of low sequence quality, MinION has been mostly used in whole-genome sequencing analyses to resolve long repeats and bridge contigs or re-sequencing genomes (Quick et al., 2014; Jain et al., 2018). The error rate of reads can be reduced from 10-15% to 1-5% by sequencing of the complementary strand (1D^2^ method) or preparing tandem repeat molecules (concatemers), but these solutions are laborious, of low sequencing depth, and hence not broadly used (Cornelis et al., 2019; Li et al., 2016; Calus et al., 2018; Volden et al., 2018). MinION has been used to generate long DNA barcodes from consensus sequences (Pomerantz et al., 2018) and to detect specific human pathogens that are easily distinguishable and well represented in reference sequence databases (Kilianski et al., 2015; Ashikawa et al., 2018). Although multiple reports claim achieving species-level taxonomic resolution (Benitez-Paez et al., 2016; Benitez-Paez and Sans, 2017; Kerkhof et al., 2017), the high error rate renders nanopore sequencing poorly suited for exploratory metabarcoding analyses of real communities. The metagenomic approach has gained popularity for identification of human pathogens to skip the PCR step and avoid associated biases (Quick et al., 2016; Schmidt et al., 2017; Votintseva et al., 2017). Recently, Bronzato Badial et al. (2018) demonstrated that plant pathogenic bacteria and viruses can be detected using MinION, whereas Hu et al. (2019) extended this to fungal pathogens of cereals in a preprint.

The main objective of this study is to develop protocols for metabarcoding-based and metagenomics-based detection of fungal plant pathogens using third-generation sequencing tools. In particular, we aim to 1) test the relative biases and shortfalls of MinION-based and Sequel-based identification of pathogens and evaluate the perspectives of these methods in pathology and ecology; and 2) optimize MinION protocols for ultra-rapid pathogen identification. We performed several HTS runs using MinION and Sequel instruments and validated the results by comparing these to Sanger sequencing, species-specific priming PCR and morphology-based assessment where relevant. We tested the third-generation HTS methods in two plant pathosystems, conifer needles and potato (*Solanum tuberosum*) leaves and tubers.

## RESULTS

### Technical features of MinION and Sequel runs

Compared with Sequel, MinION had several-fold greater initial sequencing depth, which depended on the loaded DNA content and sequencing time (Table 1). The high sequence number of MinION was reduced several-fold during the quality filtering and demultiplexing, reaching the level comparable with Sequel. Among samples, variation in sequencing depth was slightly greater in MinION (CV, 67.8%-93.4%) compared with Sequel (62.8%-64.5%). Pearson correlation coefficient of sequencing depth of samples in MinION and Sequel ranged from 0.585 in the potato data sets to 0.853 in the needle data sets, suggesting a substantial library preparation or sequencing bias in the potato amplicon pool but not in the needle sample pool.

**Table 1.**
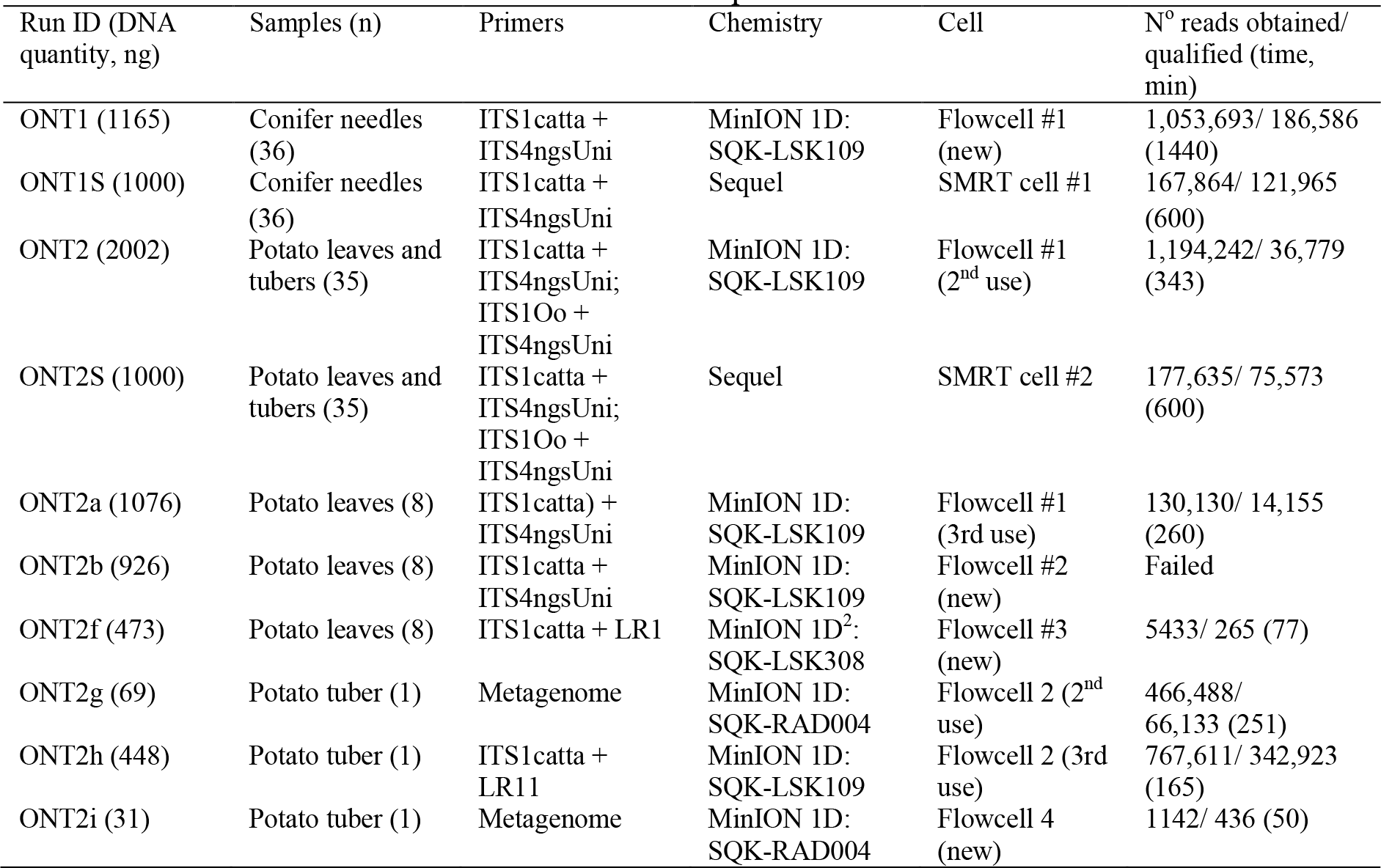
Detailed information about MinION and Sequel runs

For the MinION data sets, chimeras were detected using the reference-based method but not *de novo* method. Putatively chimeric molecules contributed 1.5%-1.8% to the mapped reads, but nearly half of these were false positives based on manual checking (cf. Hyde et al., 2013). Interestingly, nearly half of the true chimeras included parents from different samples, indicating some chimera formation during the library preparation or sequencing process in addition to PCR. Further manual inspection of demultiplexed sequences revealed that 5-8% of these are self-chimeric, i.e. 1.5-fold to 6-fold repeats of itself. In the Sequel data sets, chimeras accounted for 1.9-3.7% of reads (including 1.5-2.4% detected *de novo*), with no self-chimeric reads remaining.

Index switching rate was much greater in MinION (3.6% of reads in ONT2) than Sequel (0.14%, ONT2S). Based on positive control samples, we estimated that the error rate in Sequel is around 0.1% (corresponding to polymerase errors), but around 11-16% (depending on species) for the 1D method and 11% for the 1D^2^ method of MinION. Based on alignments of hundreds of positive control and other dominant sequence types, we noticed that errors were non-randomly distributed, i.e. occasionally there were no errors across 4-5 bases of the alignment, whereas homopolymeric sites were infested with large amounts of combined indels and substitutions (Fig. 1). Because of these non-random errors, we were able to construct consensus at 98.5-99.5% accuracy (only deletions remaining) with 100 or more reads.

**FIG 1.**
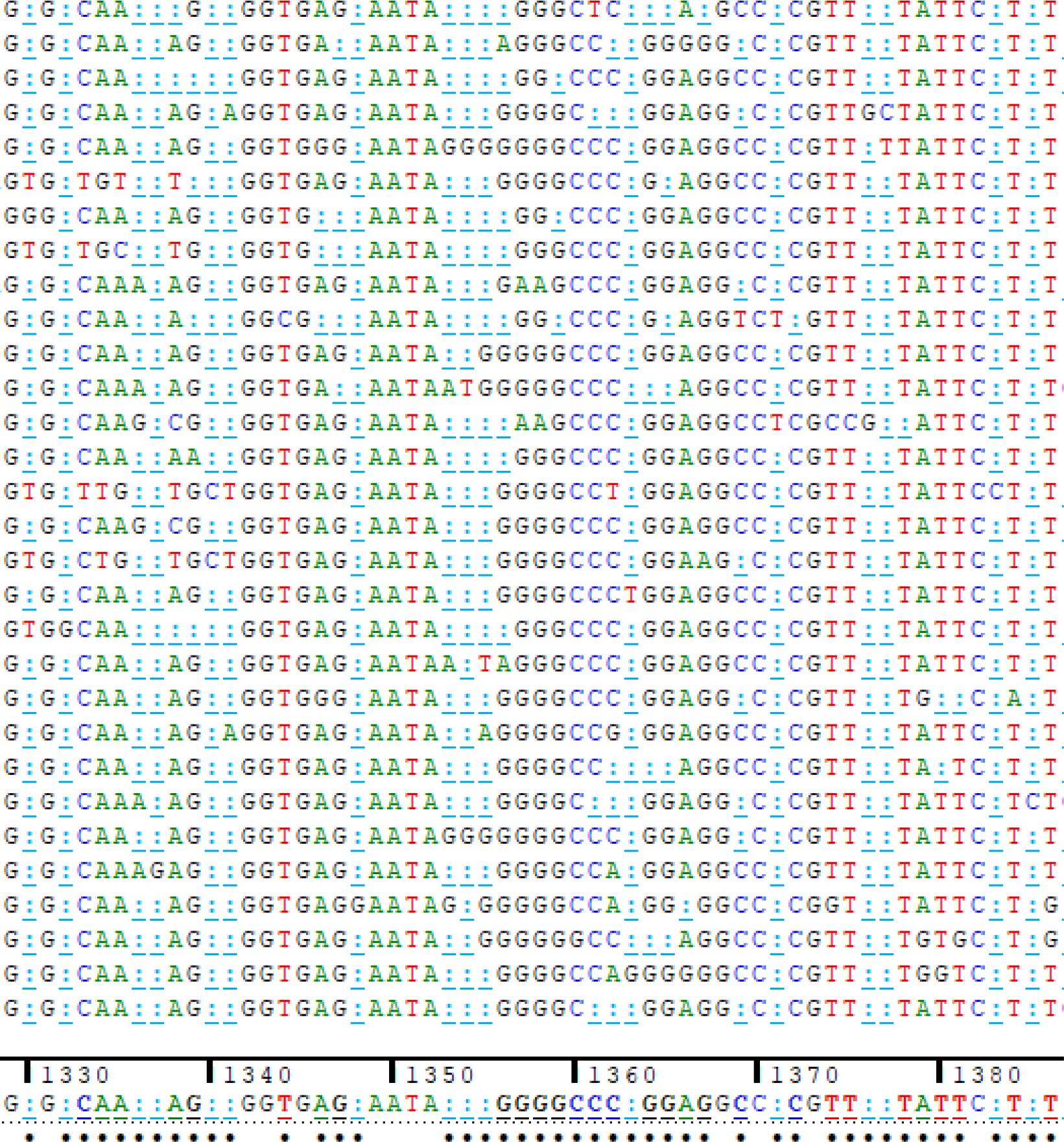
Screenshot example of multiple sequence alignment of MinION reads mapped to the contaminant *Coniothyrium* sp. using Sequencher 5.1 software (GeneCodes Corp., Ann Arbor, MI, USA). Note the error-infested double homopolymeric region (center) and a relatively accurately recorded region upstream.

All MinION runs on R9.4 flowcells from one batch (flowcells #1 and #2) were contaminated by *Coniothyrium* sp. (INSD accession JX320132), but this taxon was not observed in negative control samples, another batches (flowcell #3 and runs not reported here) or runs using R9.5 flow cells, or PacBio Sequel. At least partly because of this, the dominant fungal taxa recovered in samples differed in the MinION and Sequel runs (Tables 2, 3).

**Table 2.**
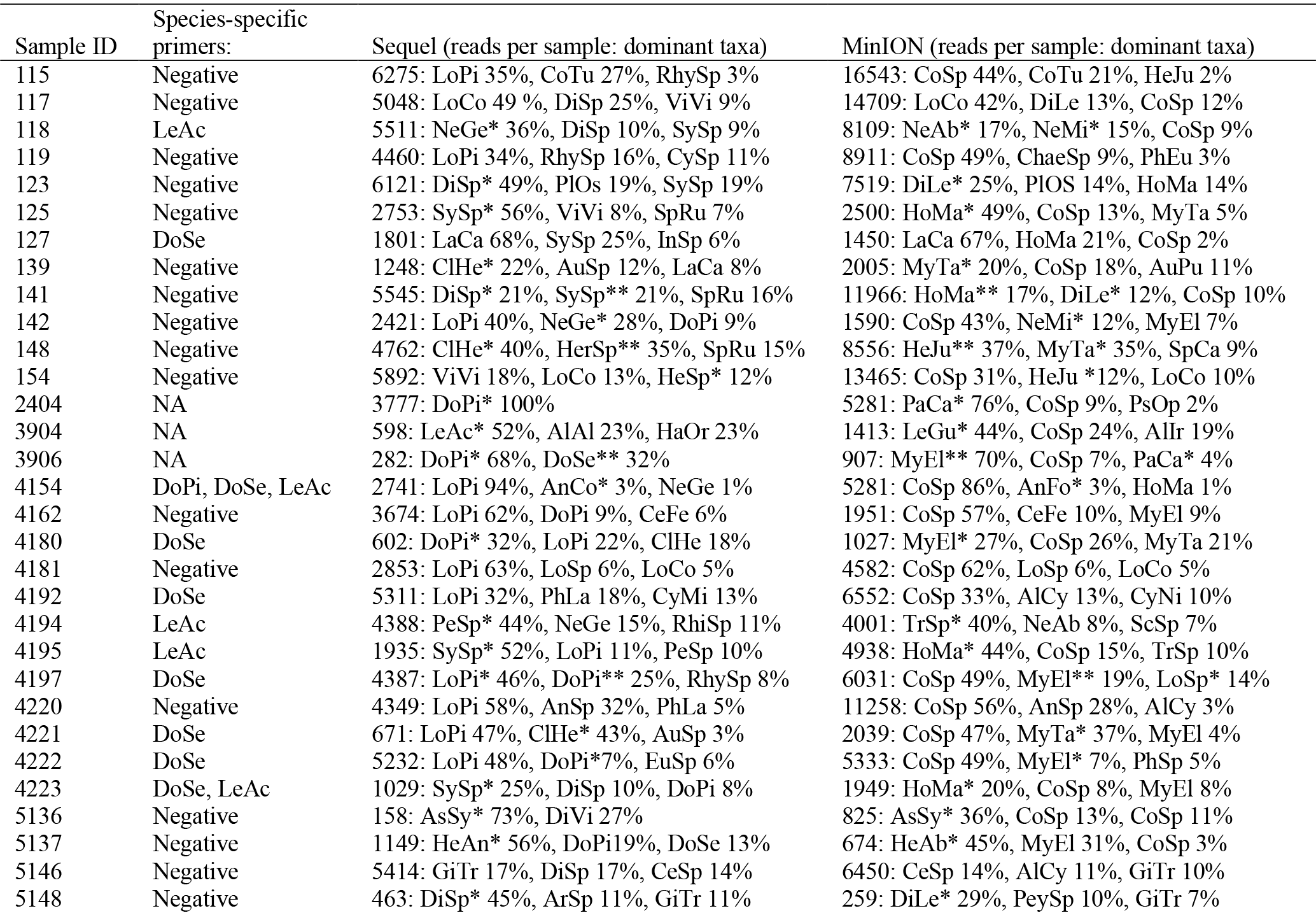
Identification of Fungi in needle samples. Numbers and percentages in Sequel and MinION columns indicate the number of fungal reads and per cent of sequences assigned to particular OTUs. Asterisks indicate taxon names that correspond to each other based on >98% sequence similarity.

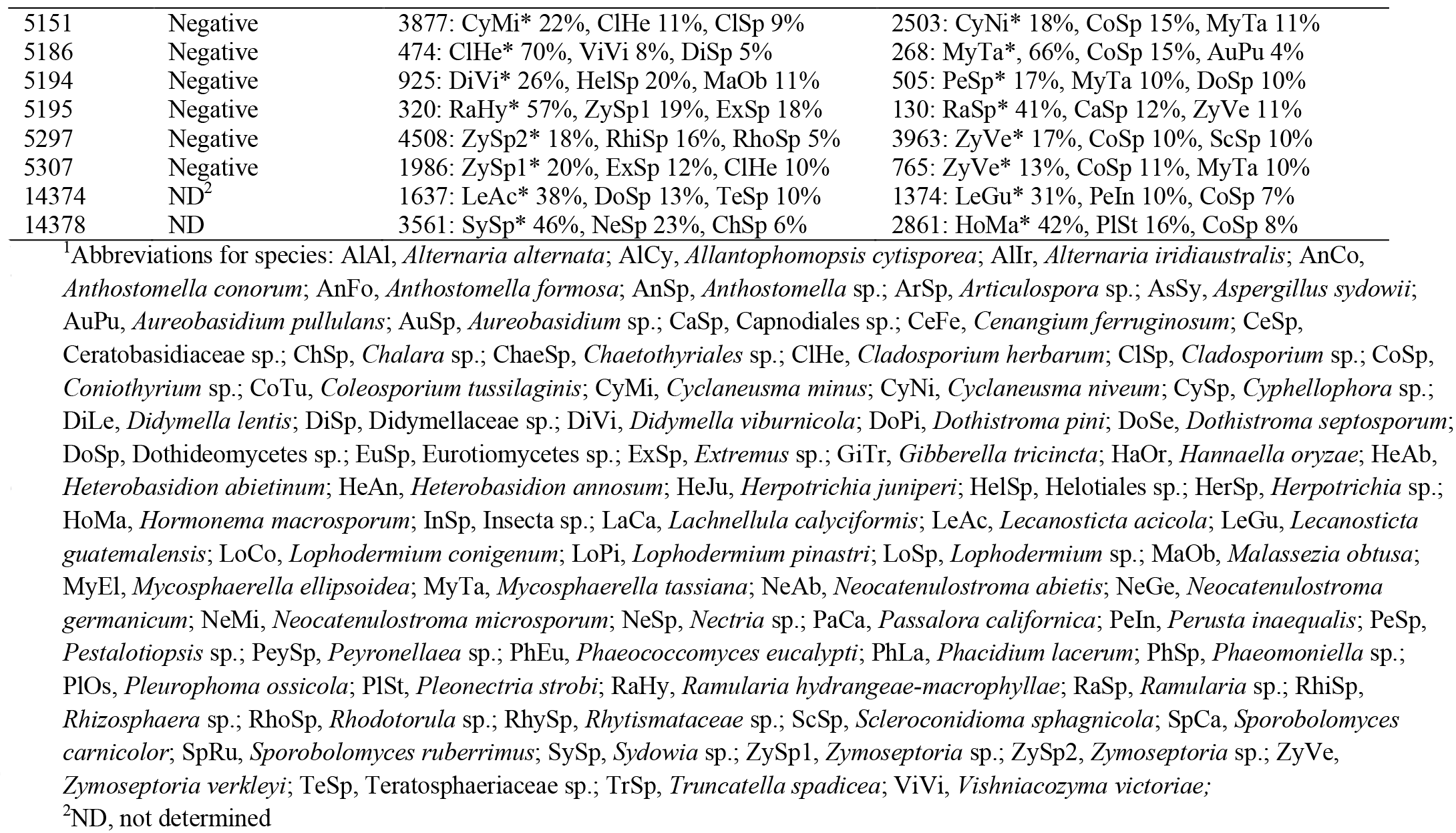

**Table 3.**
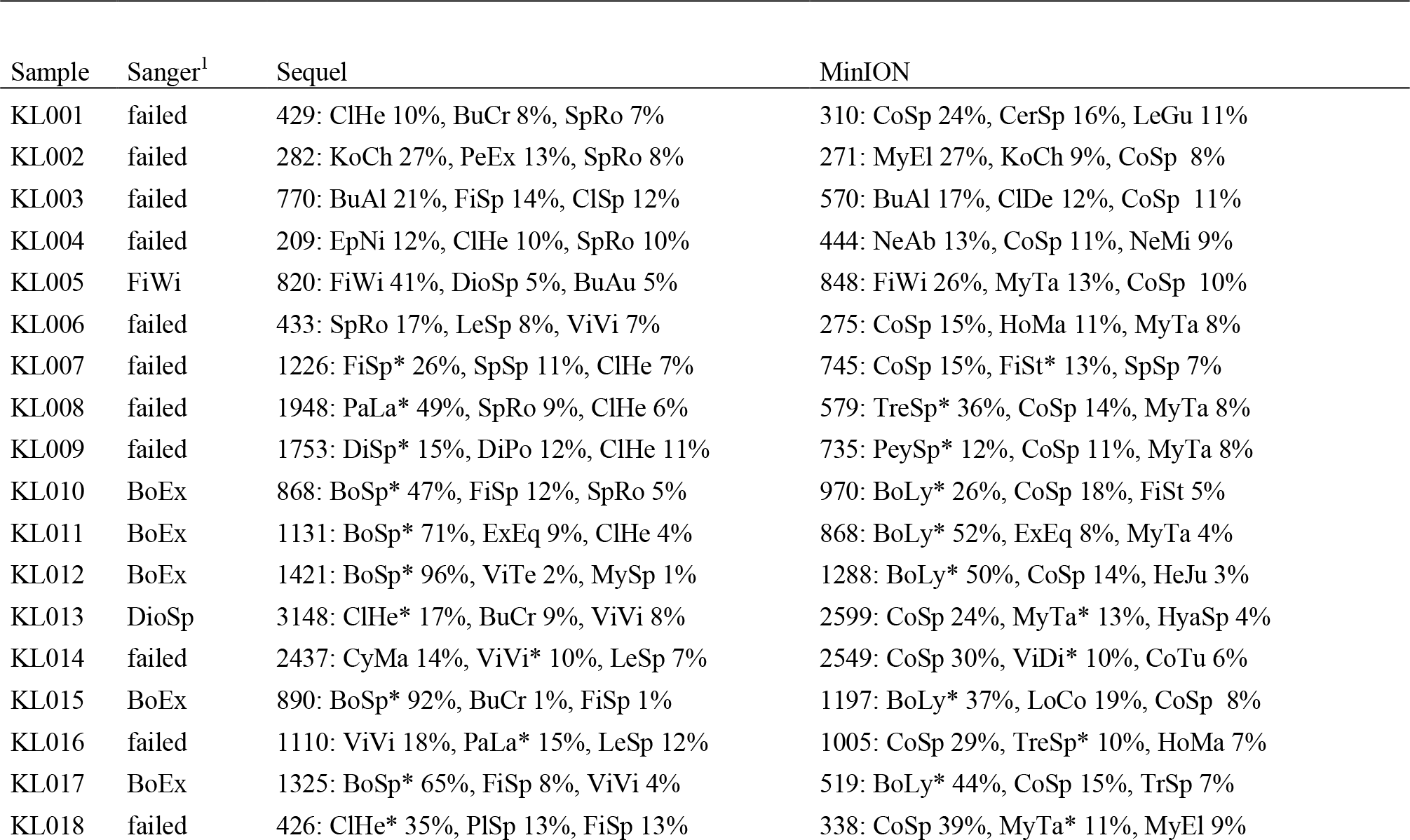
Identification of Fungi in potato samples. Numbers and percentages in Sequel and MinION columns indicate the number of fungal reads and per cent of sequences assigned to particular OTUs. Asterisks indicate taxon names that correspond to each other based on >98% sequence similarity.

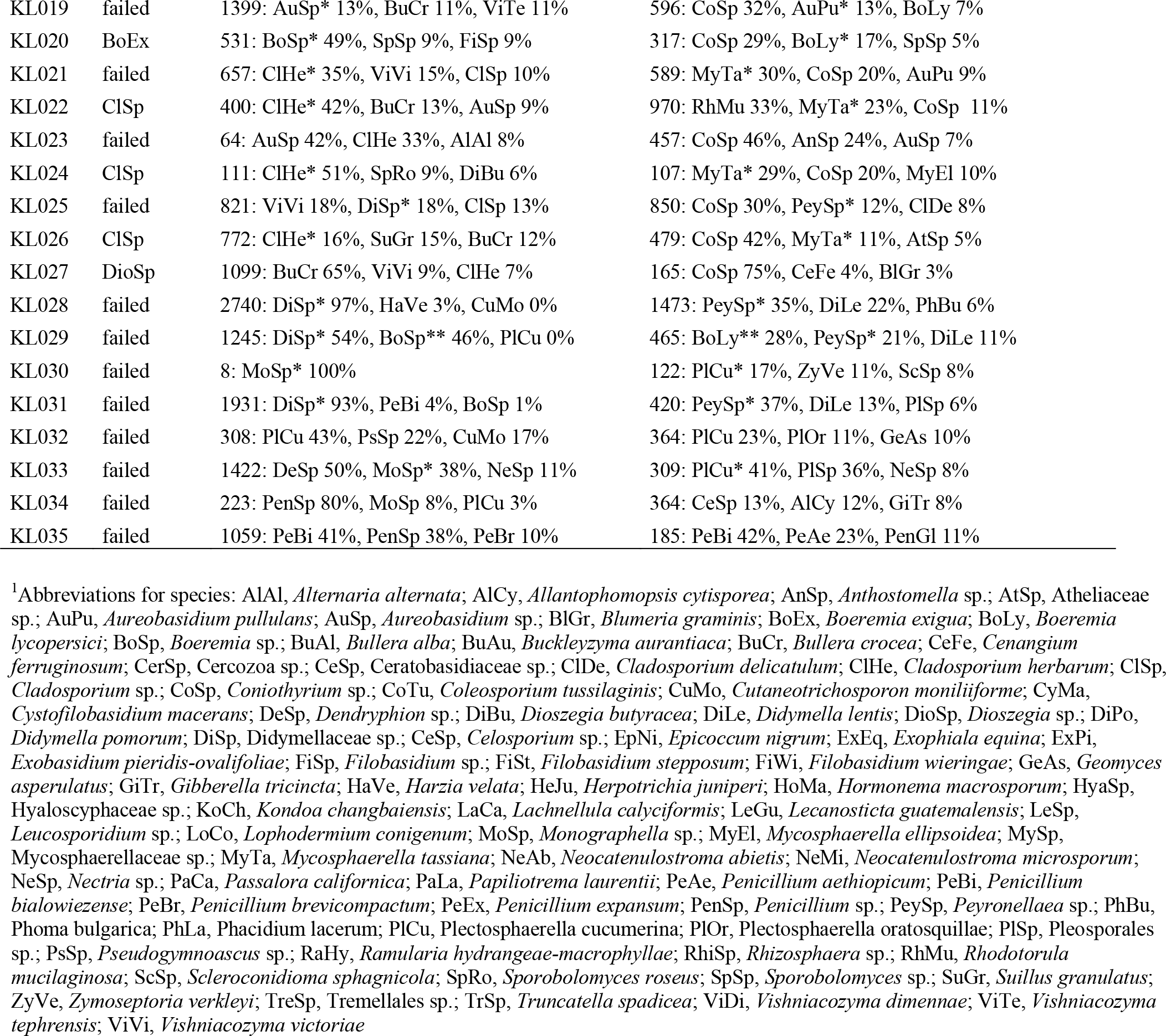

### Metabarcoding analyses of MinION and Sequel

The MinION ONT1 run included diseased and asymptomatic needle samples and pure cultures of pathogens. Of 792,748 passed reads, 189,150 (23.9%) were demultiplexed and 183,343 (23.1%) were mapped to reference sequence databases based on the quality criteria (e-value <e-40 and sequence similarity >75%). The ITS1catta forward primer amplified mostly Fungi (99.9% of identified reads). Best hits were distributed across 2483 fungal OTUs, with the well-known conifer pathogens yielding hits to 1-2 different accessions. On average, needle samples hosted 203.4±130.5 (mean±SD) OTUs. Best hits to the contaminant *Coniothyrium* sp. contributed 26.3% of all sequences on average. Of expected taxa, *Hormonema macrosporum* (6.2%), *Lophodermium conigenum* (5.0%) and *Didymella lentis* (4.3%) yielded the greatest number of hits (Fig. 2a) and all these taxa occurred in 94%-100% of needle samples.

**FIG 2.**
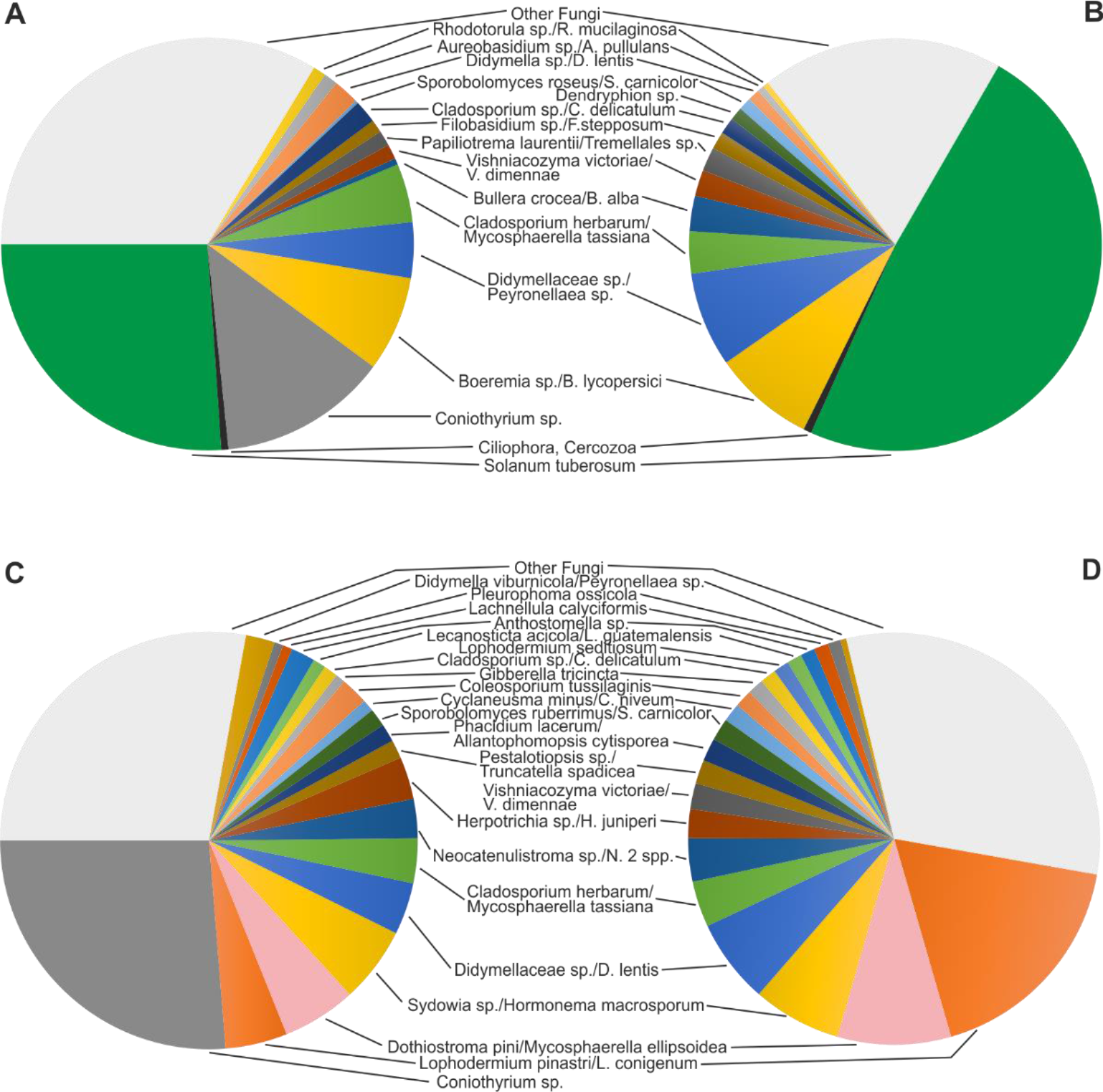
Pie diagrams demonstrating dominance of higher taxa and OTUs in a) ONT1 (needles; MinION), b) ONT1S (needles; Sequel), c) ONT2 (potato; MinION) and ONT2S runs (potato; Sequel) based on relative abundance of identified reads in the ITS1catta + ITS4ngsUni amplicons. Taxon names in Sequel and MinION data sets have been put into correspondence (/).

The ONT1S Sequel run revealed 121,965 demultiplexed reads that were clustered into 535 OTUs, all above the quality threshold. Needle samples harboured on average 51.5±41.6 OTUs, nearly four times less than in the MinION data set. Altogether 99.9% reads were ascribed to Fungi, with *Lophodermium pinastri* (17.8%), *Dothiostroma pini* (8.9%) and *Sydowia* sp. (7.0%) dominating across the entire data set (Fig. 2b). These dominant taxa occurred in 43-65% of samples.

The ONT2 MinION run recovered 255,137 passed sequences, of which 16.2% were demultiplexed and 14.4% mapped to reference database reads. Based on the distribution of pine-specific pathogens in potato samples, we estimated that 13.4% of the reads were carried over from the previous ONT1 run, in which we used the same primer and tag combinations. In the ONT2 run, these pine-specific species had a proportionally similar relative abundance when comparing across the same index combinations. The ITS1catta forward primer amplified mostly Fungi (74.2% of identified reads; Fig. 2c). Reads corresponding to potato (9 OTUs) and *Coniothyrium* sp. accounted for 26.1% and 13.2% of sequences. Of putative potato pathogens and endophytes, the ITS1catta forward primer revealed *Boeremia lycopersici* (7.5% of reads), *Mycosphaerella tassiana* (4.6%) and *Peyronellaea* sp. (4.4%) as dominants. The average richness was 81.7±43.3 OTUs per sample. The oomycete ITS1Oo primer comprised only 1.6% of all reads that were dominated by Oomycota (47.7%), other Stramenopila (19.2%), Fungi (23.6%) and Viridiplantae (9.5%). In each sample, 0-3 oomycete taxa were found and all of these occurred only once or twice (Table 4). The majority of samples produced no amplicon with the ITS1Oo primer and these samples contained no Oomycetes based on the HTS analysis.

**Table 4.**
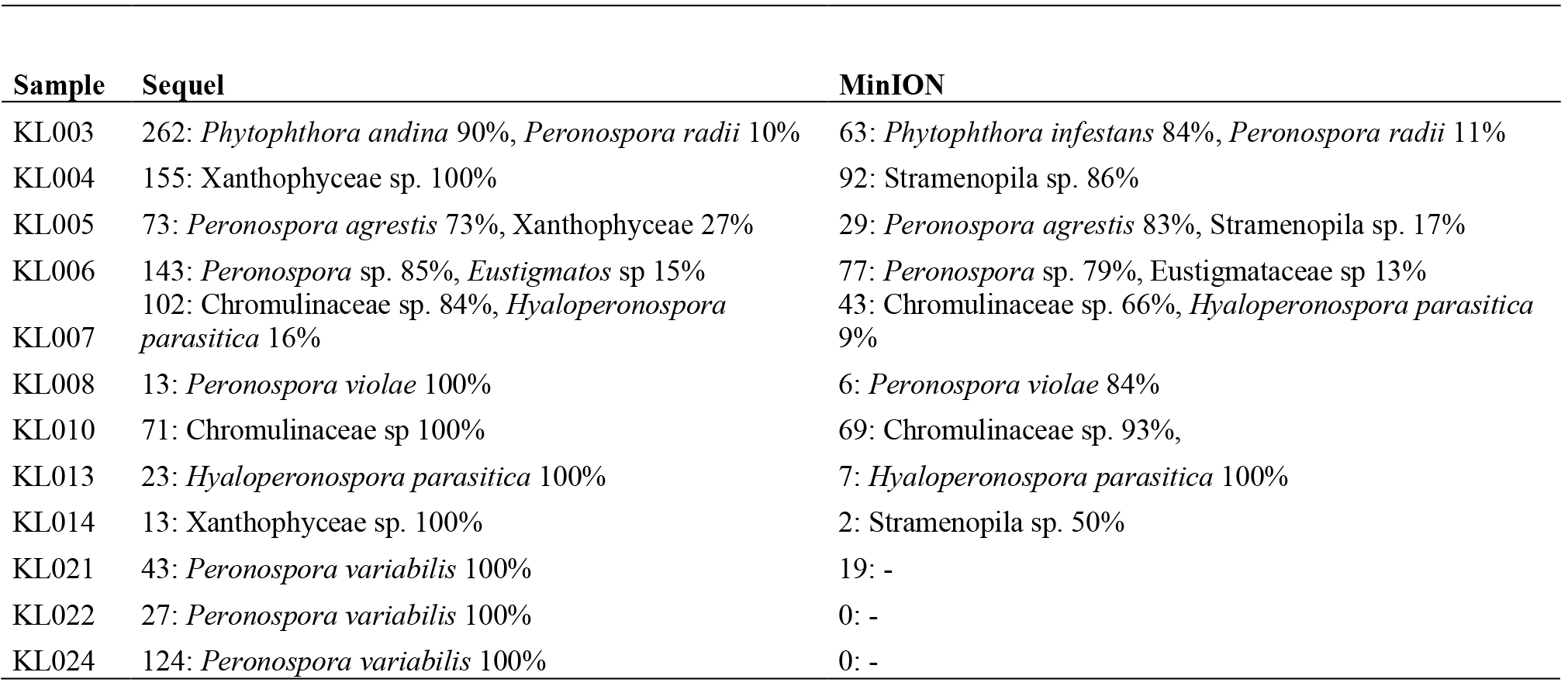
Identification of Stramenopila in potato samples based on the ITS1Oo + ITS4ngsUni primers. Numbers and percentages in Sequel and MinION columns indicate the number of all reads and per cent of sequences assigned to particular OTUs. Asterisks indicate taxon names that correspond to each other based on >98% sequence similarity. Samples with no PCR product and no sequences are excluded. Notable, plant and fungal sequences contributed on average 10% to MinION data (probably index switch artefacts from the fungal data set; not shown).

The ONT2S Sequel run revealed 75,573 demultiplexed reads that were all matched to reference sequences and separated into 308 OTUs. On average, 39.6±20.3 OTUs were recovered per sample. In the ITS1catta amplicons, Fungi, Viridiplantae, Alveolata and Rhizaria contributed to 51.0%, 48.4%, 0.5% and 0.1% of reads, respectively. All plant reads were distributed across 25 OTUs that were all assigned to potato. Six of the OTUs probably represent naturally high variation among ITS sequences of potato (based on INSD entries), whereas others represent pseudogenes or non-functional copies. These were rare to common (up to 3% of all variants) and sometimes exceeded the abundance of regular variants in individual samples. Of Fungi, the largest number of reads belonged to *Boeremia* sp. (8.0%), Hysteriaceae sp. (7.4%) and *Cladosporium herbarum* (3.3%; Fig 2.d). The ITS1Oo primer accounted for 1.4% of sequences that were mostly assigned to Oomycota (62.9%), other Stramenopila (33.9%), Viridiplantae (3.0%) and Alveolata (0.2%). This data subset yielded 0-2 OTUs of Oomycota or other Stramenopila per sample (Table 4).

Sequel and MinION recover the same dominant fungal species (excluding the contaminant) in 60% and 63% of cases in the needle and potato samples, respectively. These values increased to 78% and 83%, respectively, when considering overlap in the three best matching taxa. Inspection of the discordant samples revealed that contamination from the previous run blurred the results of the potato samples and MinION produced one to two orders of magnitude less high-quality reads matching to multiple species such as *Lophodermium pinastri, Vishniacozyma victoriae, Cystobasidium* sp. and *Dendryphion* sp. as compared with Sequel. These species had a relatively high number of homopolymers (>3-mers) per ITS sequence length compared with dominant but equally shared taxa (F_1,8_=5.79; P=0.088). The Stramenopile data subsets were in a stronger agreement in Sequel and MinION apart from the lack of *Peronospora variabilis* amongst MinION reads and hence its unsuccessful diagnosis from three potato leaf samples.

The ONT2a and ONT2b MinION runs were designed to test whether long indexes relieve the massive index switching. The ONT2a run revealed a tag switch rate of 3.8%, whereas the ONT2b run failed for unknown reasons. The potato (14 OTUs) contributed to 18.7% of reads, whereas the contaminant *Coniothyrium* species accounted for 16.7% of reads, prevailing in half of the eight potato samples. Of other fungal species, Tremellales sp. (11.5%), *Filobasidium stepposum* (7.0%) and *Mycosphaerella tassiana* (5.6%) dominated. These species were less common in these eight samples in the ONT2 run, (3.8%, 2.7% and 4.6%, respectively). Nonetheless, the same best fungal hits prevailed in 75% of the samples in the ONT2a and ONT2 runs.

The ONT2f run was intended to test suitability of the 1D^2^ method. This run recovered only 3241 1D^2^ reads. Only 29.7% of reads fell within 10% of the expected read length of ca. 3200 bases and the median read length was 954 bases. As the positive control revealed no reads, the tag switch rate could not be calculated. Of all sequences, potato (17 OTUs) accounted for 54.19% of sequences. Of Fungi (39.8%), *Taphrina populina* (6.0%), *Parastagonospora* sp. (3.8%) and *Glarea lozoyensis* (3.0%) dominated. These species were somewhat less common in the ONT2 library (0.1%, <0.1% and <0.1%, respectively). The same species were among the dominants in only 25% of samples as based on the ONT2 and ONT2f runs. It remains unknown whether these biases are related to sequencing of long amplicons or the 1D^2^ method.

### Metabarcoding vs. metagenomics approach

The ONT2g run representing a metagenome of a single diseased potato tuber sample (KL036) yielded 66,133 and 400,355 ‘passed’ and ‘failed’ sequences, respectively. The 5000 randomly selected sequences from each bin included 1325 ‘passed’ reads and 1 ‘failed’ read that met our quality standards. Altogether 37.4% of the ‘passed’ reads represented ITS sequences carried over from a previous run. After removal of these reads, the metagenomics data set was dominated by plant and bacterial reads. Best hits to *Lycopersicon esculentum* (tomato, 29.0% of reads) and seven species of *Solanum* (altogether 22.6%) collectively represented the potato. Of Bacteria, hits to *Agrobacterium tumefaciens* (10.5%), *Variovorax paradoxus* (9.7%) and *Sphingopyxis alaskensis* (3.8%) dominated. Fungal hits were less common; these to *Rhizoctonia solani* (1.6%) and *Boeremia exigua* (1.0%) prevailed. Of these taxa, *A. tumefaciens* and *V. paradoxus* are probably present given their best matches of 93% and 92% and average matches of 87% and 85% similarity, respectively, to database sequences. Conversely, *S. alaskensis, B*. *exigua* and *R. solani* are probably absent, because of their best hits reached 84%, 88% and 86%, and all hits averaged 79%, 80% and 80% similarity to reference sequences, respectively.

The ONT2h run represented a long amplicon of the same sample, recovering 342,923 ‘passed’ reads and 423,688 ‘failed’ reads. Of the randomly selected 5000 sequences, 1876 ‘passed’ reads and 1068 ‘failed’ reads met the quality threshold. The positive control used in the next to previous run accounted for 0.2% of all sequences, mostly in the ‘failed’ bin. Out of 18 most commonly hit species, the proportion of 11 differed significantly (P<0.001) among the ‘passed’ and ‘failed’ bins, indicating that reads of certain taxa are much more likely to be recorded as failed. Of the ‘passed’ sequences, matches to *Lignincola laevis* (Pleosporales, 64.3%), *Verticillium biguttatum* (Hypocreales, 5.0%) and *Thanatephorus cucumeris* (Cantharellales, 3.0%) dominated. In the fail bin, *Verticillium biguttatum* (19.9%), *L. laevis* (15.7%) and *Plectosphaerella cucumerina* (Pleosporales, 7.8%) prevailed, followed by *T. cucumeris* (6.0%). Of the dominant taxa recovered, probably only *V. biguttatum*, *T. cucumeris* and *P. cucumerina* are identified to the species level given their high maximum (>90%) and mean (>85%) blast similarity. Taxa relatively more abundant in the ‘failed’ bin tended to possess more and longer homopolymers than those in the ‘passed’ bin.

In the ONT2g metabarcoding and ONT2h metagenomics data sets derived from the same sample, none of the fungal taxa were shared. Although *R. solani* is regarded as a synonym of *T. cucumeris* (or vice versa), the isolate named as *R. solani* with available genome is probably heterospecific with the *T. cucumeris* isolate that was best matched in the amplicon data set. The *R. solani-T. cucumeris* complex has high variability in the rRNA marker genes and its taxonomy is far from settled (Veldre et al., 2013). Other fungal species common in the metabarcoding data set were absent from the metagenomics data set probably because their genome is unavailable. Several of these ascomycetes may have best matched to *Boeremia exigua* that has a genome sequence available. *B. exigua* was represented by a single read, potentially resulting from carry-over from the first run. This situation highlights limitations of the metagenomics approach when insufficient reference is available.

### Express identification

The ONT2i run intended to minimize time from sampling to diagnosis based on a single infected potato tuber sample and metagenomics approach. Using forceps, we mounted ca 20 mg of infected tissue into 2 ml Eppendorf tube containing 100 μl lysis buffer from the Phire Plant Direct PCR Kit. Based on previous optimisation for speed, we reduced the step of lysis to 15 min that included tissue disruption using bead beating (5 min at 30 Hz), brief centrifugation at 5000 g, incubation at 30 °C for 5 min and final centrifugation at 11,000 g for 1 min (the recommended protocol includes lysis without tissue disruption at room temperature >2 h). The DNA was concentrated from lysate using the FavorPrep kit following the manufacturer’s instructions except centrifugation steps for 1 min and final elution using 50 μl water (altogether 25 min). Qubit measurement revealed DNA concentration of 4.1 ng μl^−1^ (5 min). Library preparation followed the G004 protocol (50 min). The MinION run was interrupted at 1200 reads (50 min) and the 436 ‘passed’ fastq reads were analysed in PipeCraft that generated a list of 10 best hitting taxa in <5 min and revealed *T. cucumeris* as a prevalent pathogen. The parallel WIMP analysis failed because of server maintenance at the time of analysis. The entire procedure took 2 hours and 30 minutes. Notably, the sequencing process was suboptimal because of the low amount of DNA used, which resulted in <20% pores effectively used at termination of this run. Some extra time was required to finalize the results for a written report. The metagenomics reads were dominated by hits to tomato (72.7%), followed by various bacteria (6.4%), and *T. cucumeris* (5.5%), and *P. cucumerina* (3.0%) that are both known pathogens of potato. Subsequent Sanger sequencing from tuber samples with black scurf symptoms of the same diseased potato revealed *T. cucumeris* (all four subsamples) and *Pyronemataceae* sp. (50% of subsamples).

## DISCUSSION

### Use of third-generation sequencing instruments for DNA metabarcoding

Using the same amplicon pools and additional morphology-based, Sanger sequencing-based diagnosis or species-specific priming PCR, we had a unique opportunity to evaluate the relative performance and biases of MinION compared with Sequel. Nanopore sequencing revealed somewhat greater among-sample variability in sequencing depth, which may be related to library preparation, sequencing, data processing or a combination of these.

MinION suffered from a unique issue with sequence carry-over from a previous run as also noted by Cusco et al. (2018) for 6% of reads. In our study, a washing procedure with the supplier’s Wash Kite still yielded 13-37% sequence carry-over from a previous run. Furthermore, we could recover traces of a positive control used in the next to previous run at 0.2% relative abundance. It is theoretically possible that such carry-over contamination occurs on re-usable flowcells or chips of other HTS platforms as we have commonly seen it in the end of untrimmed Sanger reads.

In our analyses, MinION had an issue of contamination with a fungus matching *Coniothyrium* sp. that was not observed in Sequel run and in none of our previous data sets. INSD records indicate that this taxon is common in temperate USA. This contamination occurred in two R9.4 flowcells (#1 and #2) supplied with the MinION instrument, but not in another R9.4 batch (flowcell #3 and others not reported here) or R9.5 batch. The flowcells #1 and #2 were used over 6 months, considering several independent laboratory contamination events unlikely. Therefore, we suspect that this contamination may be related to the supplier.

Chimeric reads were common in both Sequel and MinION data. UCHIME effectively detected chimeric molecules from the Sequel data, but it performed poorly on MinION data. The error-infested reads were probably too different from each other to be recognized as chimeric. MinION data also included a substantial proportion of chimeric molecules with parents from different samples, representing a unique hybrid issue of index switching and chimera formation. A large proportion of long MinION reads represented self-chimeras that were not recognized by the chimera filtering software. This issue was common on PacBio RSII instrument (Tedersoo et al., 2018), but it was not observed in the current Sequel runs. Since MinION reads are typically mapped to reference, we estimate that the abundant chimeric molecules create virtually no bias, except those with switched tags.

Index switches during library preparation or sequencing make a strong and perhaps predominant contribution to sample ‘contamination’ (Schnell et al., 2015). The observed index switching rate of 3.6-3.8% in MinION compares poorly with that of Sequel (<0.2% in; this study) and various Illumina instruments (0.1-10%; Costello et al. 2018). The double indexes performed equally poorly, suggesting that index switches are attributable to processes in library preparation or sequencing rather than sequencing errors. At least partly, high rates of index switching spilled the dominant taxa in the deeply sequenced MinION data sets across nearly all samples and resulted in 2-fold to 4-fold greater richness per sample. Certainly, the high error rate and inaccurate mapping-based method of OTU construction contribute to this difference. The MinION-derived error-infested metagenomics and amplicon sequences may be easily mapped to various closely related species, thus elevating richness artificially. Conversely, clustering at 98% sequence similarity may be too conservative, because many pathogenic taxa differ from each other by only a few bases in the ITS region (e.g. needle pathogens *Dothistroma pini* and *D. septosporum*, see Barnes et al., 2016), and therefore several species with distinct ecology and pathology may be lumped into a single taxon (Kõljalg et al., 2013). In spite of substantial disparity in the taxon-rich MinION and Sequel fungal data subsets, these two instruments revealed high-level overlap in the species-poor oomycete data subset.

The average error rate of MinION reads was 10-15%, depending on the proportion of homopolymers in the marker gene region of particular species. Species possessing homopolymer-rich ITS markers were up to two orders of magnitude less common than in the Sequel run. This was also supported by relative prevalence of homopolymer-rich taxa in the ‘failed’ bin. We showed that this may substantially bias estimates of dominant fungal and perhaps oomycete taxa in specific samples and overall.

We observed discrimination against longer reads when sequencing potato amplicons using the 1D^2^ approach (ONT2f), which confirms a previous report (Cusco et al., 2018). Preferential recovery of shorter reads seems to be inherent to both PCR and all sequencing instruments including Sequel (Tedersoo et al., 2018, 2019; Nilsson et al., 2019).

Using the same amplicon pools, Sequel and MinION revealed contrasting results in metabarcoding of conifer needle and potato samples. The results of Sequel were generally consistent with morphology-based and species-specific priming PCR diagnosis (needle samples) and Sanger sequencing results (potato samples), but failed to differentiate the closely related needle pathogens *D. pini* and *D. septosporum.* Apart from the species of *Coniothyrium* contaminating the MinION data sets, the two platforms revealed different taxa (by names and INSD accessions) prevailing in the same samples. In some cases, these taxa are considered synonyms (e.g. *M. tassiana* and *C. herbarum*), but mostly these belong to closely related sister taxa that share the UNITE Species Hypothesis at 2% level as confirmed by manual comparisons of best-matching reads. In nearly 20% of occasions, inconsistencies between MinION and Sequel were attributable to the poor recovery of taxa with abundant homopolymers in ITS sequences by MinION. Possibilities to solve this include lowering of the initial phred score or including the ‘failed’ reads in analyses.

### Rapid molecular diagnostics

We tested both metagenomics- and amplicon-based approaches of nanopore sequencing for rapid identification of pathogens. Most taxa recovered in the metagenome run were rare in the amplicon data set and vice versa. Although we detected severe biases in MinION amplicon sequencing, we believe that amplicon-based analyses are more accurate and that reference bias accounts for much of the difference; i.e., in metagenomics analyses, species with available reference genomes have much greater chance of accumulating hits compared with species with no available genomes. In our analyses, this is illustrated by misidentification of potato as a tomato. Mapping reads of an ascomycete pathogen to genomes of several others, as in our study, is likely to remain cryptic. A solution may be sufficiently deep metagenomics sequencing to secure coverage of mitochondrial or ITS marker genes that are in multiple copies per genome. Because genomes of most bacterial and fungal human pathogens have available genome sequences, nanopore metagenomics-based identification may be better suited to medical samples, but this situation is likely to improve very soon in plant pathology.

We demonstrate that accurate molecular identification from sample collection through DNA extraction, concentration, library preparation, sequencing, bioinformatics and taxonomic interpretation may take as little as two and half hours using MinION sequencing in the metagenome mode. This is less than the four hours reported for identification of bacterial human pathogens from urine, which requires no specific DNA extraction (Schmidt et al., 2017), and one working day as commonly reported in multiple studies (e.g. Quick et al., 2016; Votintseva et al., 2017; Charalampous et al., 2018). The protocols for plant and animal tissues can be potentially optimised to 90 minutes in cases where DNA is easily extractable in high quantity and purity; library preparation and sequencing process can be limited to 10-20 minutes each, when using rapid library kits and halting efficient sequencing runs after a critical number of reads (Votintseva et al., 2017). These express diagnostics rates of MinION cannot be beaten by instruments of other HTS platforms that require several hours for library preparation and at least one day for sequencing (except 6 hours for Ion Torrent; Reuter et al., 2015).

However, there is a clear trade-off between overall analysis time and data reliability in nanopore sequencing. Shortened DNA extraction protocols tend to yield lower quality and quantity of DNA, whereas culled incubation and centrifugation steps in library preparation may result in dilute and poorly indexed libraries overrepresented by short fragments. Although we successfully identified potato pathogens from a library 13-fold more dilute than recommended, it may be useful to add a certain amount of so-called carrier DNA to secure preparation of high-quality libraries (cf. Mojarro et al., 2018). Sequencing time is almost linearly related to sequencing depth and thus quality of consensus reads and genome or taxonomic coverage. Sample preparation, bioinformatics and interpretation processes take longer for multiplexed samples, which may be necessary to reduce the overall analytical costs of 500-800 EUR per sample to ca 100 EUR per sample using Oxford Nanopore’s commercial multiplexing kits or to 2-3 EUR per sample by using custom multiplexing methods of indexed primers. For example, analytical costs for the ONT1 and ONT2 runs were roughly 10 EUR per sample considering triple use of flowcells. To reduce the chances of carryover of previous molecules, contamination-aware indexes (different indexes across runs) could be used.

### Technical and analytical issues

Although several authors report species-level identification of bacterial taxa in complex communities using MinION (e.g. Shin et al., 2016; Kerkhof et al., 2017), these interpretations are not convincing, because mapping of sequences with 10-25% errors to reference reads of high similarity is inaccurate (see above). We demonstrate that even when using the relatively rapidly evolving fungal ITS region and reference sequences differing from each other by at least 2%, positive control samples and plant material yield multiple OTUs, sometimes recovering strongest hits to different genera. Conversely, species absent from databases are mapped to one or more closest species, which may provide wrong taxonomic implications. This is of particular importance for molecular diagnostics, necessitating inclusion of marker genes of all potential target species in the reference database to prevent incorrect interpretation. This issue is particularly pressing for long rRNA amplicons and single-copy genes. The metagenomics approach requires a comprehensive database of genomes of all potential target organisms, which strongly depends on whole genome sequencing initiatives. Exome compilations are suboptimal, because much of the eukaryotic DNA is non-coding. Besides target organisms, metagenomics databases also require inclusion of potential contaminants such as specific interacting taxa (e.g. potato) and human, and various bacteria that contribute much to the metagenomic DNA.

A major concern with novel sequencing techniques is the paucity of reports on analytical shortfalls, and nature of artefacts, which may partly be derived from the lack of controls or inappropriate sampling design (Pontefract et al., 2018). The MinION has been used for five years, but so far there is very little information about analytical errors and biases, and very few authors mention about checking chimeras, index switching artefacts or unsuccessful runs (but see White et al., 2017 Cusco et al., 2018; Xu et al., 2018). The virtual lack of constructive (self)criticism echoes a misleading message about the multi-purpose, non-problematic use of the method, serving the interests of the manufacturer and journal editors in an unjustified manner. Users of MinION, many of which have no prior experience with other HTS techniques and related problems, heavily struggle to squeeze reasonable data out of the device. There are thousands of academic users, but only a few hundred papers out. For example, our team purchased MinION instruments with extra troubleshooting service; yet, the company responded only to technical questions but not to troubleshooting about index switching, contamination and sequence carry-over. It should be the responsibility of researchers and editors to include such problematic issues and solutions in publications to prevent the research community and specialists of governmental and private institutions from wasting countless time (re)falling into the same analytical holes.

### Perspectives of third-generation sequencing technologies

Both MinION and Sequel are rapidly evolving in terms of read length and base calling accuracy. At the moment, Sequel seems to be the best choice for metabarcoding regular-size (600-1000 bp) and long (up to 3 kb) amplicons (Cline et al., 2015; Kyaschenko et al., 2017; Tedersoo et al., 2018; Nilsson et al., 2019; Tedersoo et al., 2019) and for barcoding ultra-long markers (up to 7 kb; Wurzbacher et al., 2019). The forthcoming M8 chemistry will reduce the costs of PacBio sequencing roughly two-fold. Declining costs, greater throughput, read length and quality continue to make Sequel more attractive for metabarcoding and seek supporters from metagenome and metatranscriptome researchers (Rard, 2018). However, it may be hard to convince users of Illumina MiSeq to switch to another platform and adopt alternative bioinformatics workflows.

Use of MinION for metabarcoding looks relatively less promising considering the current state-of-the-art technology with unacceptably high error rates. The error rates should be reduced to <0.1%, i.e. to the level of Illumina and circular consensus of Sequel for use in routine metabarcoding. Several methods of tandem repeat (concatemer) sequencing enable to reduce error rates to 1-3% (Li et al., 2016; Calus et al., 2018; Volden et al., 2018). Double-barcoding of each size-selected RNA or DNA molecule followed by generation of consensus sequences yields quality improvements comparable to tandem repeat sequencing (Karst et al., 2018), but it would require ultra-high sequencing depth to reach 1% error rate and to be able to multiplex over several biological samples. Combining these methods may facilitate reducing error rates towards the critical threshold.

For regular barcoding, the third-generation sequencing tools offer great promise in situations where their throughput and read length are much superior compared to double-pass Sanger sequencing, i.e. for barcodes >1000 bases and multiplexing hundreds of samples to secure cost-efficiency (Hebert et al., 2018; Srivathsan et al., 2018). Sanger sequencing handles poorly the alleles or copies of markers with read length polymorphism, which is common in non-coding regions of eukaryotes. The third-generation HTS technologies are able to recover various alleles as well as pseudogenes (Cornelis et al., 2019) from mixed or contaminated samples by sequencing single DNA molecules. Sequel and MinION are capable of handling DNA amplicons of >7000 bases, requiring generation of consensus reads for reliable results (Wurzbacher et al., 2019). Although we could reach 99.5% accuracy with over 100 reads, Pomerantz et al. (2018) estimated that 100 MinION reads suffice for principally error-free generation of barcodes for animals using sequences from a complementary strand. For PacBio, a single read may be enough for reads around 2000 bp, but three or more may be needed for longer fragments and to average over PCR errors.

Unlike Sequel and other HTS technologies, MinION is well-suited to rapid diagnostics of pathogens and invasive species especially in groups that are well-known and well-referenced in public sequence databases. Besides detection of pathogenic species, MinION has a potential to recover antibiotic resistance genes and pathogenesis-related genes as well as single mutations in the metagenomics mode (Bradley et al., 2015; Cornelis et al., 2019). By using multiplex amplicons or metagenomics/genomics approach, it will be possible to detect harmful organisms and their specific pathogenicity-related genomic features in less than one day (Schmidt et al., 2017). Besides enabling to trace taxon/gene exchange between different habitats (Bahram et al., 2018), this approach has important implications for improving diagnosis and implementing countermeasures, e.g. releasing biocontrol agents, spraying biocides or arranging quarantine.

Nanopore-based detection methods are flexible for incorporating additional options such as recording epigenetic modifications (Jain et al., 2016; Simpson et al., 2017) and primary structure of RNA (Garalde et al., 2018), proteins (Perez-Restrepo et al., 2018) and polysaccharides (Karawdeniya et al., 2018). Alternative nanopore-based DNA sequencing methods are also evolving (Goto et al., 2018). The potential of different nanopores to record various biomolecules indicates great promise of nanopore-based diagnostics in the future.

### Conclusion

We demonstrate that the MinION device is well-suited for rapid and accurate diagnosis of pathogens, which may take as little as 150 minutes from sample preparation (including DNA extraction, library preparation, sequencing, bioinformatics and data interpretation). However, care should be taken to secure profound reference sequence data to avoid misdiagnosis. Amplicon-based diagnostics takes longer time, but is more accurate if genomes of potential pathogens are unavailable. For whole-community metabarcoding, Sequel performs much better than MinION in terms of data quality and analytical biases. Although development of tandem repeat sequencing and read consensus sequencing have been developed for MinION, their error rate of 1-3% is still insufficient for exploratory metabarcoding analyses of biodiversity, but this may change in the coming years.

## MATERIALS AND METHODS

### Sample preparation

The potato subset includes 27 samples of leaves and 10 samples of tubers with symptoms of disease (Table 5). We also included a DNA sample of two Australian Tuberaceae species as a positive control. The conifer system includes 36 distinct needle samples from eight species of *Pinus* and two species of *Picea* that represent material with symptoms of needle blight or no symptoms. The samples of natural, planted or recently imported trees were collected in Estonia in 2011-2018 (Table 6). We included a cultured isolate of *D. pini* (146889), *D. septosporum* (150931) and closely related *Lecanosticta acicola* (150943) as positive controls. Unequal mixture of DNA from these cultures served as a simple mock community. In both systems, we included a negative control sample.

**Table 5.**
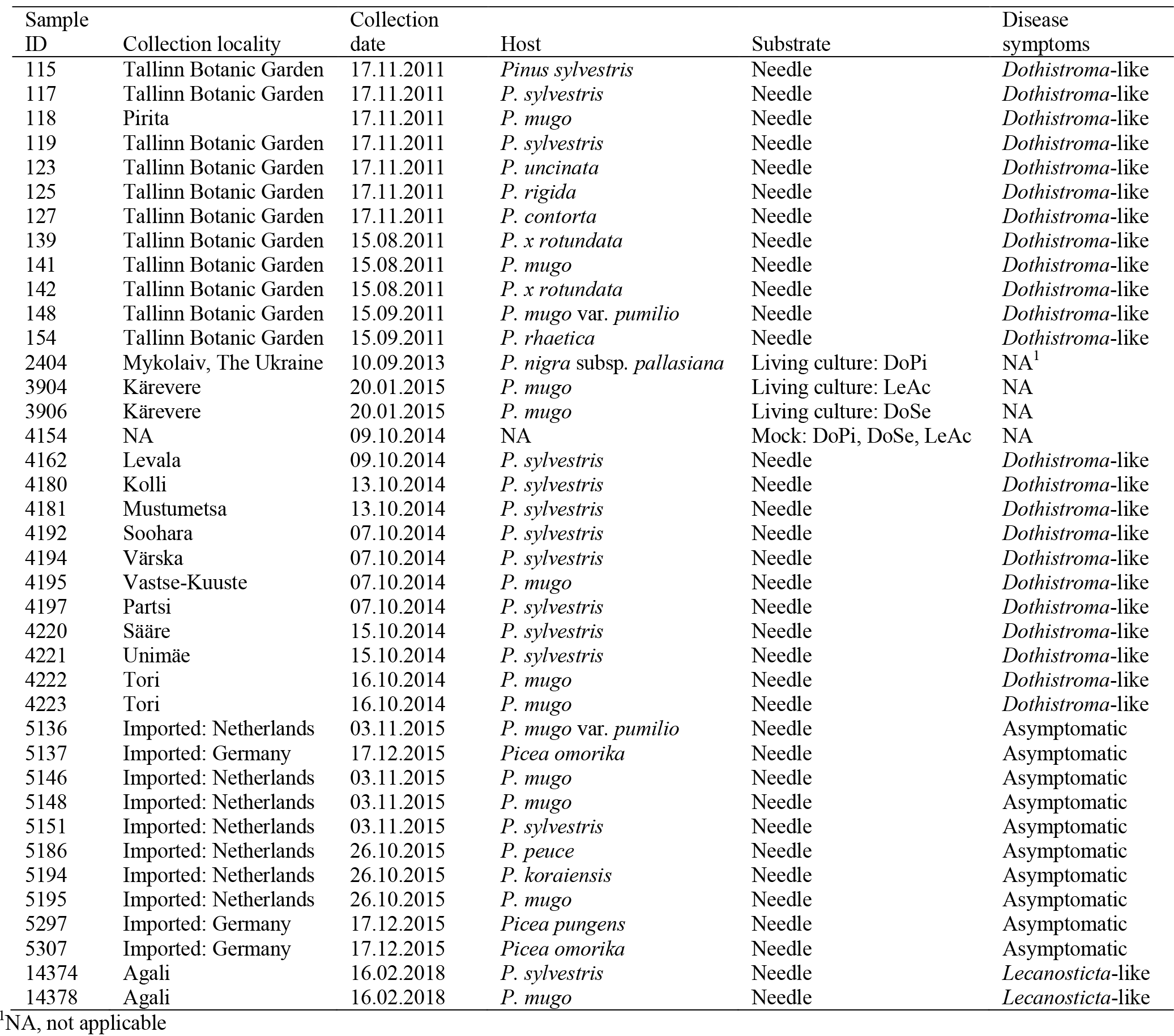
Details of needle samples.

**Table 6.**
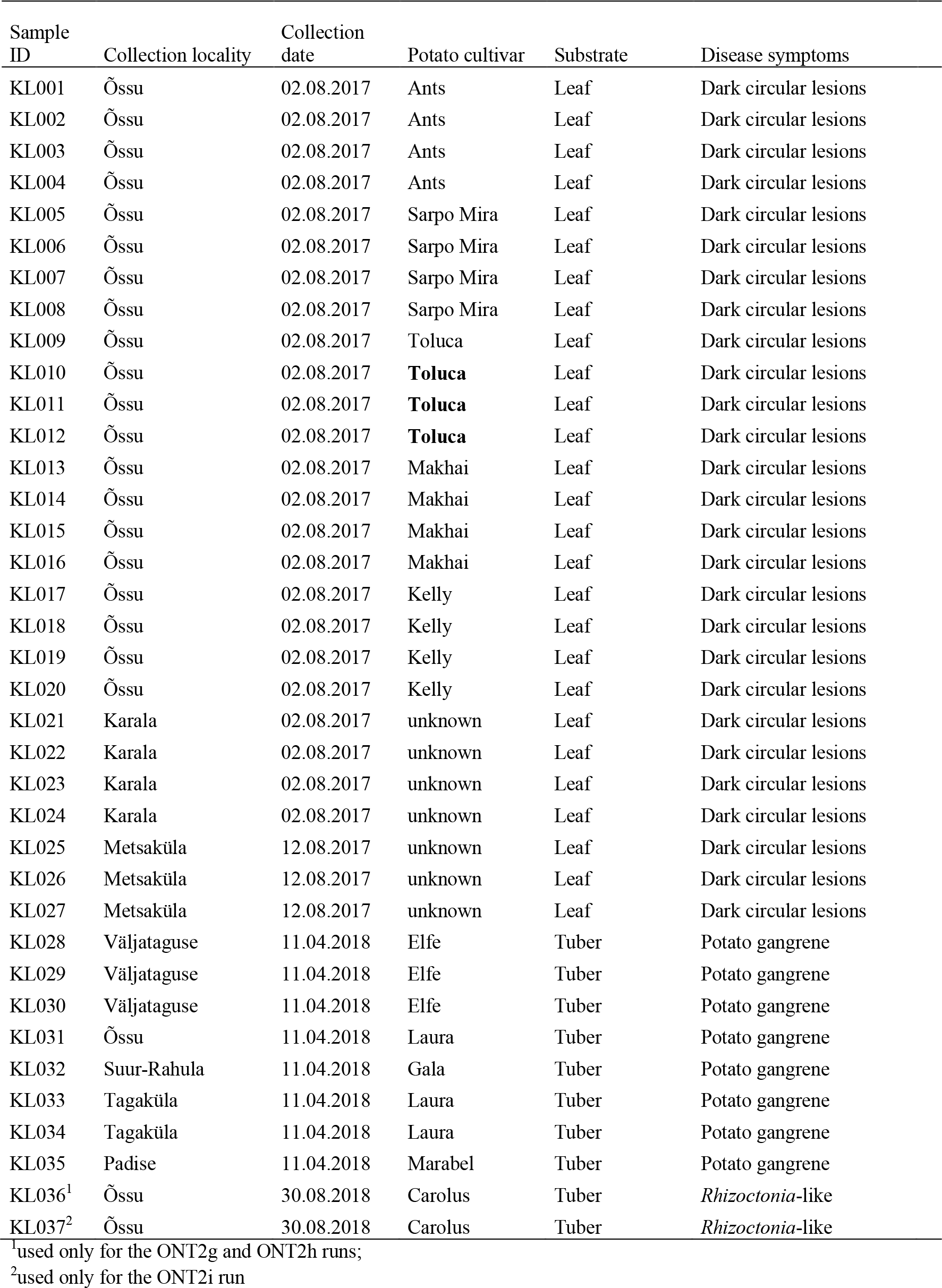
Details of potato samples.

In needle samples, DNA from 0.2 g plant material and fungal cultures was extracted using the Thermo Scientific GeneJET Genomic DNA Purification Kit (Thermo Fisher Scientific, EU). In potato samples, DNA from 0.1 g diseased fresh leaf tissue was extracted with a lysis buffer (0.8 M Tris-HCl, 0.2 M (NH_4_)_2_SO_4_, 0.2% w/v Tween-20; Solis BioDyne, Tartu, Estonia). For some additional analyses using potato samples, we also used Phire Plant Direct PCR Kit (Thermo Scentific, Waltham, MA, USA).

### Molecular identification

Needle samples were screened for specific pathogenic fungi *Dothiostroma pini, D. septosporum* and *Lecanosticta acicola*, using species-specific primer pairs following the developers’ protocols (Ioos et al., 2010). Potato samples were amplified using the ITS1F + ITS4 primer pair (White et al., 1990; Gardes & Bruns, 1993) and sequenced using the ITS5 primer (White et al., 1990).

For the metabarcoding approach, we used a forward primer ITS1catta (5’-ACCWGCGGARGGATCATTA-3’; Tedersoo et al. submt.) and a reverse primer ITS4ngsUni (Tedersoo & Lindahl, 2016) to be able to selectively amplify fungal DNA and simultaneously avoid the 18S rRNA gene intron bias. Located in the terminus of the 18S rRNA gene, the ITS1catta primer covers nearly all Ascomycota and Basidiomycota as well as selected groups of ‘zygomycetes’ and ‘early diverging lineages’, but discriminates against plants and most other eukaryote groups (incl. fungal taxa Mortierellomycota and Tulasnellaceae) in the last position. To specifically detect Oomycota, we used the ITS1Oo primer (Riit et al., 2016, 2018) in combination with the ITS4ngsUni primer for the potato data set. Forward primers were tagged with one of the 12-base Golay indexes with at least four differences to any other index (Tedersoo et al., 2018). Because of issues in data recovery, we also amplified a subset of eight potato samples (KL001-KL008) using ITS1catta and ITS4ngsUni primers in which the forward primer was equipped with tandem repeat barcode of double length (securing at least 8 base difference) to increase resolution among samples. Because the 1D^2^ nanopore sequencing method requires DNA fragments of >2kb, we amplified these potato samples (>3 kb amplicons) using the indexed ITS1catta primer combined with the LR14 primer (Vilgalys & Hester, 1990). For comparing the metabarcoding approach to metagenomics method, we used ITS1catta in combination with the LR11 primer (Vilgalys & Hester, 1990) that yielded stronger amplicons compared with LR14. We used negative and positive controls as above.

PCR was performed in 25 μl volume comprising 0.5 μl each of the tagged primer (20 μM), 5 μl HOT FIREPol Blend Master Mix (Solis Biodyne), 1 μl DNA extract and 18 μl ddH_2_0. Thermocycling conditions included an initial 15 min denaturation at 95 °C, 30 cycles of 30 sec denaturation, 30 sec annealing at 55 °C and 60 sec elongation at 72 °C, and a final 10 min elongation before hold at 4 °C. The number of cycles was increased to 35 or 38 for some samples to yield a visible amplicon on 1% agarose gel. For the ITS1Oo + ITS4ngsUni primer combination, 40 PCR cycles at 50 °C annealing was used to secure greater product recovery. The two replicate amplicons were pooled, checked on a gel, and mixed with amplicons of other samples in roughly equal proportions based on visual estimates of band size.

### Third-generation sequencing

The mixed amplicons of potato and those of needles were separately split into library preparation for Sequel and MinION. The two PacBio libraries were sequenced on a Sequel instrument using the same SMRT cell (SMRT cell 1M, v2; Sequel Polymerase v2.1, sequencing chemistry v2.1., loading by diffusion, movie time 600 min, pre-extension time 30 min). PacBio CCS reads (minPasses=3, MinAccuracy=0.9) were generated using SMRT Link v 6.0.0.47841 (Pacific Biosciences). Subsequent bioinformatics were performed using PipeCraft 1.0 (Anslan et al., 2017) that included steps of demultiplexing (2 mismatches to primer and 1 mismatch to tag), extraction of the ITS region (ITSx: default options; Bengtsson-Palme et al., 2013), chimera removal (UCHIME: de novo and reference-based methods combined; Edgar et al., 2011; and additional filtering option „remove primer artefacts” that removes chimeric sequences were the full length primer is found somewhere in the middle of the read), clustering (UPARSE: 98% sequence similarity threshold; Edgar, 2013), taxonomic identification (BLASTn: e-value=0.001, word size=7, reward=1, penalty=−1, gap open=1, gap extend=2; Camacho et al., 2009) against UNITE 7.2 (Kõljalg et al., 2013) reference database. We used the criteria of blast e-value <e-40 and sequence similarity >75% for the kingdom level identification, and e-value <e-50 for phylum and class-level identification.

For the MinION instrument, amplicon library preparation was performed using the 1D amplicon/cDNA by Ligation (SQK-LSK109) kit (Oxford Nanopore) using R9.4 flowcells following manufacturer’s instructions. For long fragments, we also used the 1D^2^ sequencing of genomic DNA (SQK-LSK308) kit on R9.5 flowcell, following the producer’s protocols. Flowcells were used 1-3 times, being cleaned once or twice using the Wash Kit (EXP-WSH002; Oxford Nanopore). Sequencing was performed in the laboratory at room temperature, connecting the MinION device to a plugged-in, internet-connected laptop computer with four processors and 32 GB RAM. For base calling in MinKnow 3.1.19 software (Oxford Nanopore), we used the default phred score = 11, which placed the reads into ‘passed’ and ‘failed’ bins. The ‘passed’ fasta-formatted reads (additionally ‘failed’ reads in some analyses) were further subjected to bioinformatics using PipeCraft and WIMP (Juul et al., 2015) in parallel. The options in PipeCraft included demultiplexing of metabarcoding reads allowing no mismatches to the barcode, followed by blastn search using default settings. The sequencing adaptors were removed by a custom script.

Given the poor overall sequence quality, traditional OTU-based approaches are not applicable to the MinION data; therefore, we mapped reads based on their best matches to database sequences in the UNITE reference database, following previous nanopore sequencing studies (Benitez-Paez et al., 2016; Kerkhof et al., 2017). Limitations of this approach are outlined in the Discussion.

To maximize the speed of pathogen diagnosis, we used a metagenomics-based approach with the MinION instrument. For this, we concentrated the genomic DNA of select potato samples using FavorPrep Gel/PCR Purification kit (Favorgen Biotech Corp., Vienna, Austria). For library preparation, the Rapid Sequencing kit (SQK-RAD004) and Rapid Barcoding Sequencing kit (SQK-RBK004) were used following manufacturer’s instructions. Base calling was performed as described above. Both ‘passed’ and ‘failed’ sequences were used in further bioinformatics analyses as implemented in the Pipecraft. Taxonomic reference libraries included UNITE 7.2 and SILVA 132 (Quast et al., 2013), and INSD for extracted rRNA gene reads and other reads separately. The UNITE database was merged with a database of oomycetes created based on ITS sequences in INSD. Using this reference, chimera checking was performed using Uchime on demultiplexed reads. Specific information about the amount of initial material, sequencing time and sequencing runs is given in Table 1.

Sanger sequences of potato samples have been deposited in the UNITE database (https://unite.ut.ee/; accessions UDBxxxxx-UDBxxxxx). Raw sequence data of MinION and Sequel are available from the PlutoF digital repository. Sample-by-OTU tables used in these analyses are given in supplementary material (Tables S1 and S2).

## ACKNOWLEDGEMENTS

We thank V. Kisand for constructive comments on the manuscript and A. Tooming-Klunderud for running PacBio sequencing. Financial contribution was provided by the Estonian Science Foundation (grants PUT1399, PUT1317, PSG136, IUT21‐04, IUT 36-2, MOBERC13, ECOLCHANGE). Author contributions: KL, RD and LT designed the study; RK, KL, KA and RD provided material; RP and KL performed HTS analyses; MB, SA and LT analyzed data; LT wrote the paper with input from all co-authors.

## APPENDIX

## Additional File 1

**Table S1** Sample-by-OTU table of metabarcoding studies of conifer needles.

**Table S2** Sample-by-OTU table of metabarcoding studies of potato tissues.

